# Neural network control of focal position during time-lapse microscopy of cells

**DOI:** 10.1101/233940

**Authors:** Ling Wei, Elijah Roberts

**Author notes:** Correspondence to: Elijah Roberts, Department of Biophysics, Johns Hopkins University, Jenkins Hall 110, 3400 N Charles St, Baltimore, MD 21218.

## Abstract

Live-cell microscopy is quickly becoming an indispensable technique for studying the dynamics of cellular processes. Maintaining the specimen in focus during image acquisition is crucial for high-throughput applications, especially for long experiments or when a large sample is being continuously scanned. Automated focus control methods are often expensive, imperfect, or ill-adapted to a specific application and are a bottleneck for widespread adoption of high-throughput, live-cell imaging. Here, we demonstrate a neural network approach for automatically maintaining focus during bright-field microscopy. Z-stacks of yeast cells growing in a microfluidic device were collected and used to train a convolutional neural network to classify images according to their z-position. We studied the effect on prediction accuracy of the various hyperparameters of the neural network, including downsampling, batch size, and z-bin resolution. The network was able to predict the z-position of an image with ±1 *μ*m accuracy, outperforming human annotators. Finally, we used our neural network to control microscope focus in real-time during a 24 hour growth experiment. The method robustly maintained the correct focal position compensating for 40 *μ*m of focal drift and was insensitive to changes in the field of view. Only ~100 annotated z-stacks were required to train the network making our method quite practical for custom autofocus applications.

## 1 Introduction

Biologists now routinely use live-cell imaging to monitor the dynamics of the cell’s state, to track real-time biochemical processes *in vivo,* and to read out cellular decision-making processes from time-lapse microscopy images [1, 2, 3]. From these experiments, it has become apparent that cell populations inevitably display significant cell-to-cell variability in cellular state, as quantified by protein and RNA expression levels [4, 5, 6]. Of particular interest, populations are often observed to contain cells exhibiting rare phenotypes, i.e., small subpopulations with a distinct state [7, 8, 9]. Studying the dynamics of these small populations is indispensable for understanding the probabilistic principles behind how cells make transitions to these rare, but stable, phenotypic states [10, 11, 12, 13, 14]. The challenge in live-cell imaging, then, becomes to record large populations such that rare phenotypes are sufficiently sampled for statistical analysis, perhaps requiring 10^5^–10^6^ total cells per experiment.

With advanced optical instruments and computer-controlled stages, time-lapse microscopy is a viable method for capturing high-throughput data regarding cellular phenotypes [15,16]. Additionally, data-intensive computing methodologies are now available that can analyze the large volumes of imaging data that are generated [17]. One challenge to bringing high-throughput microscopy into every biology lab, though, is its performance on less sophisticated equipment. Keeping the cells of interest in focus during time-lapse experiments and/or continuous movement is difficult [18]. Due to thermal drift, uneven slide surfaces, diverse cell sizes, and cell motion in 3D space, there is not a single focal plane that will accommodate the entire experiment [19, 20]. In order to maintain image sharpness throughout the experiment, focus need to be corrected before each image is acquired. Manual control of focal position is not practical for experiments running for hours or when specimens are in continuous motion. Instead, automated focusing is needed to establish the best focus in the absence of human intervention [21].

Autofocus methods can be broadly classified into two categories: active and passive autofocus. Active autofocus uses knowledge of the physical characteristics of the system to obtain defocus information, and then correct the defocus accordingly [22]. Typically, electromagnetic waves, such as a laser or infrared light, are applied to monitor the distance between object of interest and lens [20]. This approach is able to provide real-time correction, however, what it actually measures is the distance between the reflective surface and lens, rather than the specimen itself [19]. Therefore, any variations in thickness of the sample or coverslip can introduce error in determining the correct focal plane [21]. Moreover, active autofocus methods require calibration before focal length can be determined, which makes it impractical for many image acquisition processes [23].

In contrast, passive autofocus is based on image analysis, and thus requires little knowledge of the specific imaging system [22, 24]. In passive autofocus, a predefined focal reference is first determined, typically by taking a series of images at multiple z-positions on both sides of best focus of the sample. The images are then processed and high frequency contents or edge information is extracted from the z-stacks to establish the focal position [20]. Since passive autofocus is based on the information of the sample being imaged, it is generally more reliable than active autofocus [21]. Nevertheless, passive autofocus suffers from limitations that are intrinsically hard to overcome. Low image intensity or low contrast may prevent it from being accurately analyzed [20]. In addition, because multiple images at different z-positions need to be acquired and processed in order to determine the relative focus, it takes longer to refocus the sample. Therefore, passive autofocus is not ideal for tracking changing objects or for high-throughput imaging acquisitions.

As a long standing topic over the years, autofocus in high-throughput microscopy has been approached by a number of methods, most of which provide satisfying solutions under well-defined circumstances [25]. Still, these methods are designed for a single type of imaging mode, and thus not applicable to other imaging systems. In recent years, deep learning has become a promising approach for inference across various fields, such as speech recognition, visual object recognition, object detection, drug discovery, and genomics [26]. It allows computational models with multiple processing layers to learn features of data by extracting information from multiple levels [26]. Specifically, deep convolutional neural networks (CNNs) are especially helpful in visual object recognition and object detection, for instance, annotation of cellular cryo-electron tomograms, classification of cancer tissues using hyperspectral imaging, cell segmentation in fluorescence microscopy images, and autofocusing in digital holographic microscopy [27, 28, 29, 30, 31]. CNNs are essentially neural networks that employ the convolution operation as one of its layers. Typically, a CNN consists of a feature extraction layer that extracts features from input data and a classification layer that classifies the feature map, the weights of which are determined through a training process [26]. The feature extraction layer consists of pairs of convolution and pooling layers in tandem, where the convolution layer performs convolution operation on input data, and the pooling layer reduces the dimension of the input data. The output of CNNs is generated from the classification layer, which in most cases employs an ordinary multi-class classification neural network.

In this paper we demonstrate a deep learning approach to use convolutional neural networks as an autofocus method for light microscopy. A benefit of the proposed method is that the z-position of an image can be determined solely from the information in the image itself, without the necessity of using other z-plane images. Thus, real-time control of focus can be realized, which allows high-throughput data acquisition from time-lapse and fast-tracking microscopy. Using the proposed method we achieved accurate control of the focal position in bright-field microscopy of yeast cells, and the method was shown to be robust against noise, optical artifacts, and features other than cells.

## 2 Results

### 2.1 Autofocus neural network structure, training, and testing

To infer the focal position of a yeast cell culture during live-cell microscopy, we built and trained a convolutional neural network (CNN) using z-stacks of yeast cells growing in a microfluidic device. Our network is shown in Figure 1a and uses a variation on a well-studied CNN architecture for image classification [32, 33]. The CNN consists of two convolution blocks followed by two fully connected layers. Each convolution block consists of a convolution, an activation, and a max-pooling layer. The first convolution layer maps image regions to 32 features with a kernel size of 10×10×1 with a stride of 1 and zero padding to preserve the image size. A rectified linear unit (ReLU) activation function is applied to the convolution output to provide a nonlinearity. Finally, a 5×5 max-pooling layer with a stride of 5×5 downsamples the spatial domain of the features. The second convolution layer maps the downsampled features from the first block to 64 larger-scale features with a kernel size of 10×10×32 and, similarly, ReLU activation and 5×5 max-pooling are performed afterwards.

**Figure 1:**
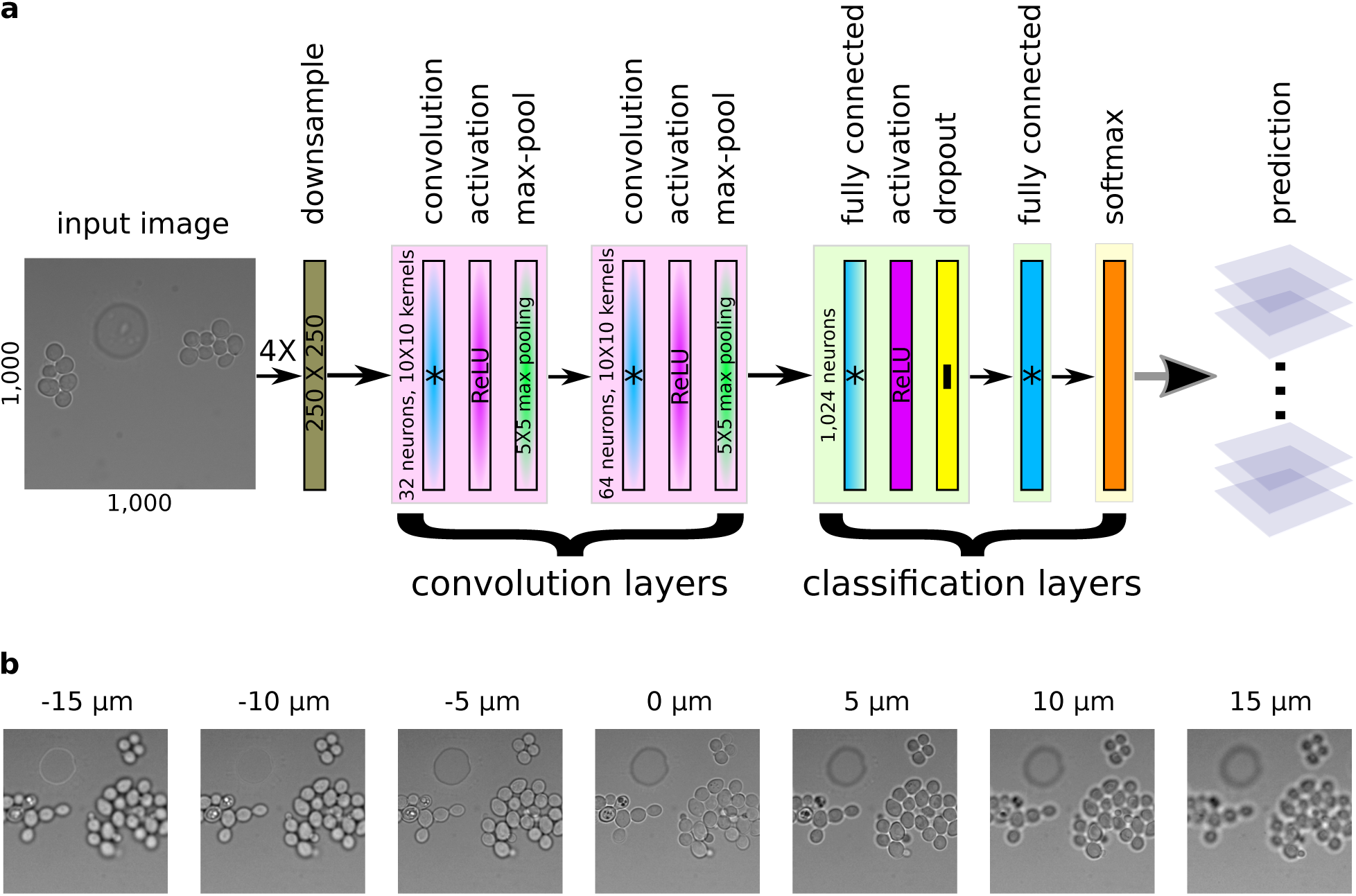
Autofocus convolutional neural network. (a) Architecture of the convolutional neural network (CNN). Images of 1000x1000 pixels in size are downsampled before entering the neural network. The CNN consists of two convolution blocks and two fully connected layers. (b) Sample images from a z-stack. Each z-stack is composed of 31 images in steps of Δ*z* = 1 *μ*m, including images below and above the focal plane (*z* = 0). The complete z-stack is shown in Figure S1.

The first fully connected layer maps the feature output from the second convolution block to 1024 classification neurons. It is followed by ReLU activation and a dropout layer to minimize overfitting [34]. The dropout probability was set to 0.5. The last fully connected layer maps the output of neurons from the previous layer to a variable number of neurons that predict the discretized z-position of the image, using one neuron for each possible z-bin. Finally, a softmax layer gives the z-position probabilities. For reference, a model using 4X downsampling and 41 z-bins has 6.8 million parameters.

We collected 431 z-stacks of yeast cells (see Figure 1b) from five independent growth experiments for use in training and testing the neural network (see Methods). Each z-stack spanned ~30 *μ*m of focal distance using 31 images collected ~1 *μ*m apart in the z dimension. We manually curated each z-stack and recorded the z-slice that was considered in focus, usually near image 15. We randomly partitioned the z-stacks into training (80%) and testing (20%) data sets.

To train the network, we followed the typical procedure for image classification [32]. Briefly, we used stochastic gradient descent on mini-batches containing 10–50 images to optimize the multinomial logistic regression. We initialized all weights in the neural network with random Gaussian distributed values with mean 0.0 and standard deviation 0.1, truncated at two standard deviations. All biases were initially set to 0.1. To generate each image in the mini-batch, we randomly selected an image from the training data set and then extracted a random 1000×1000 pixel region from it. We then randomly flipped and rotated this image to augment the data set and to compensate for global illumination effects. Finally, the image was downsampled by 2–10X. To perform the optimization we used the Adam algorithm [35] with a training rate of 10^−4^, *β*_1_ = 0.9, and *β*_2_ = 0.999. We took the cross entropy between the neural network output and the annotated focal position as the loss function. Each CNN model was trained for 100,000 steps, which was roughly 20X coverage of the training data.

To evaluate each model we used the testing data set. We extracted the center 1000×1000 pixel region from each image in the testing data set and downsampled it to be consistent with the neural network model being tested. The processed images were evaluated by the trained neural network and the inferred focal position of the image was taken to be the center of the z-bin with the highest predicted probability. We then calculated the distribution of errors between the annotated and predicted focal positions for the entire testing data set. Overall, the CNN was able to infer the focal position of an image remarkably well. Using the best model, half of all the test images were predicted within ±1 *μ*m of the annotated z-position. We then studied the effect on prediction accuracy of the various hyperparameters of the CNN model.

### 2.2 The effect of downsampling on inference accuracy

To investigate the effect of both feature size and resolution on CNN accuracy, we evaluated three different image downsampling factors: 2X, 4X, and 10X (see Figure 2a-c). In the full resolution image, a large yeast cell with a diameter of 6 *μ*m would be ~100 pixels across. For 2X downsampling this corresponds to a diameter of 50/10/2 pixels in the various convolution layers, 25/5/1 for 4X downsampling, and 10/2/0.4 for 10X downsampling. We trained a new CNN model using data downsampled with each of these three values.

**Figure 2:**
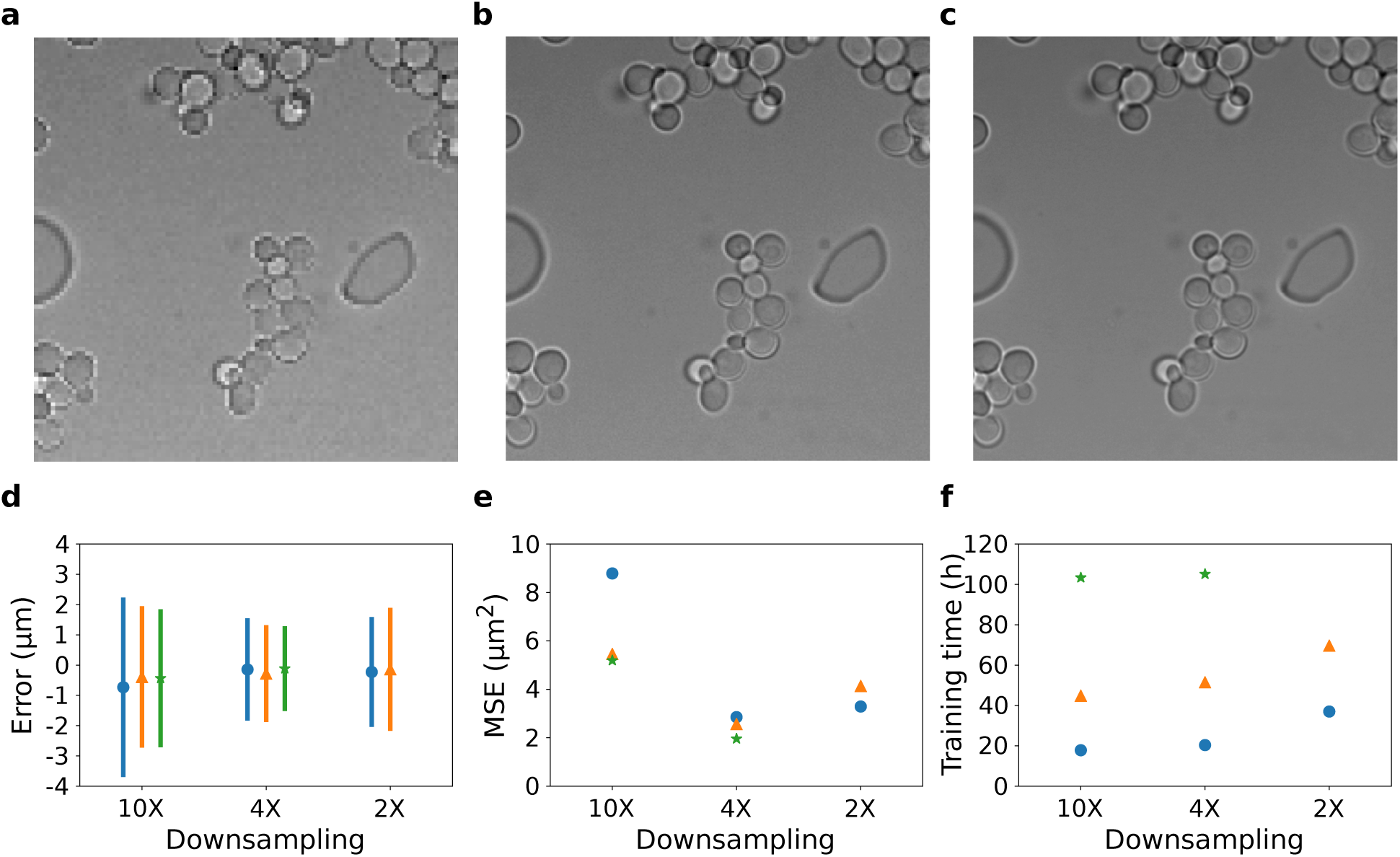
The effect downsampling on CNN accuracy. (a-c) Images downsampled by (a) 10X, (b) 4X and (c) 2X from an original image of 1000x1000 pixels in size. (d) Error of CNNs trained with different downsampling and batch sizes. The error is defined as the difference between the inferred and annotated z-position. Symbols give the mean error and bars are ±1 standard deviation. Blue dots, orange triangles, and green stars correspond to batch sizes of 10, 25 (22 for 2X downsampling) and 50, respectively. (e) Mean squared error (MSE) with symbols and colors as described in (d). (f) Time taken to train each CNN.

We analyzed the full test data set using each of the trained neural network models. The errors for models trained with different downsampling factors are plotted in Figure 2d. The mean error stays relatively consistent when varying downsampling. However, the error distribution is wider for downsampling of 10X and 2X than downsampling of 4X. Figure 2e shows the mean squared error for the inference. The model trained with a downsampling of 4X exhibited a lower mean squared error than with the models trained with 10X and 2X. Notably, in the 4X downsampling model a typical cell with an initial diameter of 60-100 pixels will be represented by a spatial domain of around ~1 square pixel in the final convolution layer. It appears that this represents a reasonable choice when deciding on the construction of a CNN for classifying cells. In the 10X downsampling model, cells occupy less than a single pixel so multiple neighboring cells become averaged before the classification layer is applied. In the 2X downsampling model cells would occupy a roughly 2×2 pixel area, which also appears to be less optimal, perhaps because of the increase in the number of parameters that need to be optimized.

We also tested the effect of varying batch size on the models. For the models trained with downsampling of 10X and 4X, the mean squared error increases with decreasing batch size, which also can be seen from the larger distributions in Figure 2d. With larger batch sizes the gradient is calculated from more training data, which helps improve training performance in general. Even though models trained with larger batch size give better results, the training time increases significantly with batch size, as shown in Figure 2f. It took more than 100 hours to train our models with a batch size of 50. In addition, for a specific batch size the training time also increases with reduced downsampling.

### 2.3 Inference accuracy depends upon the z-bin size

A key advance in our study is the classification of images into z-position bins. Thus, the number and spacing of the bins may have a large effect on the accuracy of our method. To investigate whether and how bin size affects the z-position prediction of our CNN, we tested various bin sizes ranging from 1 *μ*m to 5 *μ*m. All of the bins spanned the range −20 *μ*m to +20 *μ*m, covering the entire focal distance of our z-stacks. Thus, the output of the final classification layer ranged from 9–41 categories. We assumed that the inferred z-position was the center of the predicted z-bin.

Figure 3a shows the probability of a particular focal position inference given a label, for the highest resolution model with a bin size of 1 *μ*m (41 categories). Near the center of the focal plane, the correct bin has the highest probability in all cases with a lower amount of probability appearing in the ±1 bins. Near the edges of the z-stack the probability becomes more spread out as these images become more difficult to distinguish. The model has the highest accuracy when the cells are near the focal plane. Figure 3b shows the probability map for the lowest resolution model with a bin size of 5 *μ*m (9 categories). Here, the model is able to accurately infer the single correct bin, except when the actual z-position approaches the boundary between two bins. But the localization error is overall greater because the width of the bins is greater.

**Figure 3:**
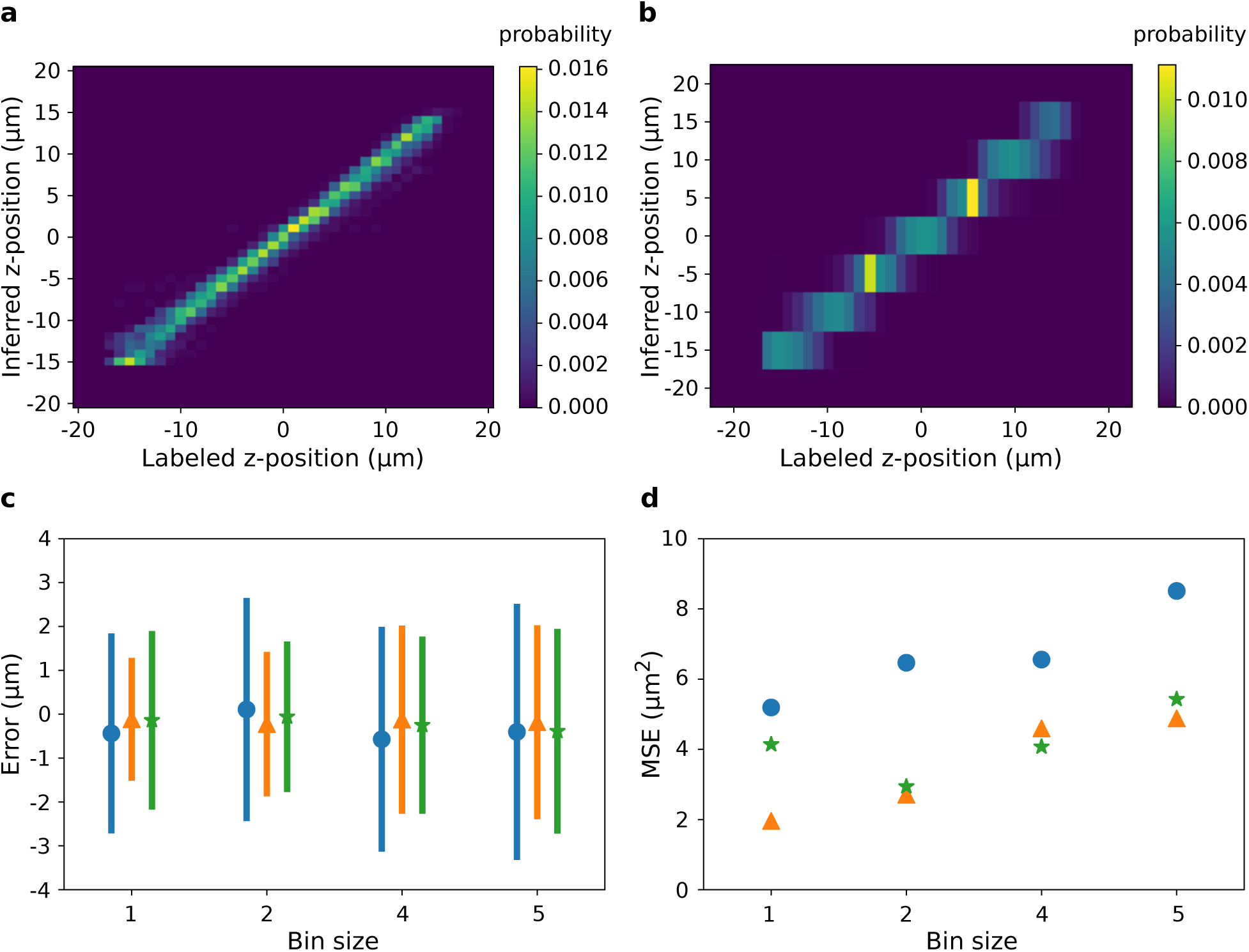
Accuracy of CNNs with different z-bin sizes. (a-b) The probability of inferring a z-position given the actual z-position using a CNN with a bin size of (a) 1 and (b) 5. (c) Error as a function of bin size. Symbols give the mean error and bars are ±1 standard deviation. Blue dots, orange triangles, and green stars correspond to downsampling of 10X, 4X and 2X, respectively. A batch size of 50 was used for 10X and 4X downsampling, and batch size of 22 was used for 2X downsampling. (d) Mean squared error (MSE) vs. bin size.

Figure 3c-d show the inference error by bin size. The width of the error distribution and the mean squared error both increase monotonically with bin size. If the z-bin resolution dropped below the information content of the images we would expect there to be a minimum in the mean squared error. However, for our z-stack resolution even a one-to-one mapping between z-slices and bins produces good accuracy. It appears that there are sufficient differences between z-slices separated by a single step to accurately differentiate them. With higher resolution z-stacks further improvement in the positional inference could likely be achieved by the model.

### 2.4 The effect of training data set size on CNN performance

In the previous sections, we used our complete training data set consisting of 345 z-stacks to train the CNN. However, it would be useful to know the minimum number of z-stacks needed to achieve good focal position inference. Thus, we trained additional CNNs using only a subset of the training data in order to investigate how the size of training data set affects the accuracy of trained CNNs. We randomly selected 50, 100, 150, 200, 250 and 300 z-stacks from the complete training data set to train the new models.

The prediction errors of these trained neural networks are shown in Figure 4a. The width of the error distribution drops at ~100 z-stacks. Similarly, the mean squared error (Figure 4b) also drops sharply at ~100 z-stacks and then continues to slowly decrease as more z-stacks are used. Therefore, while the CNN trained with the full training data set outperforms the networks trained with smaller data sets, it is possible to train a network to high accuracy with only 100 z-stacks. The collection and annotation of unique z-stacks represents a significant fraction of the total effort to build an autofocus CNN model. Collecting data incrementally until the model’s accuracy converges, rather than collecting a large data set up front, appears to be a reasonable strategy.

**Figure 4:**
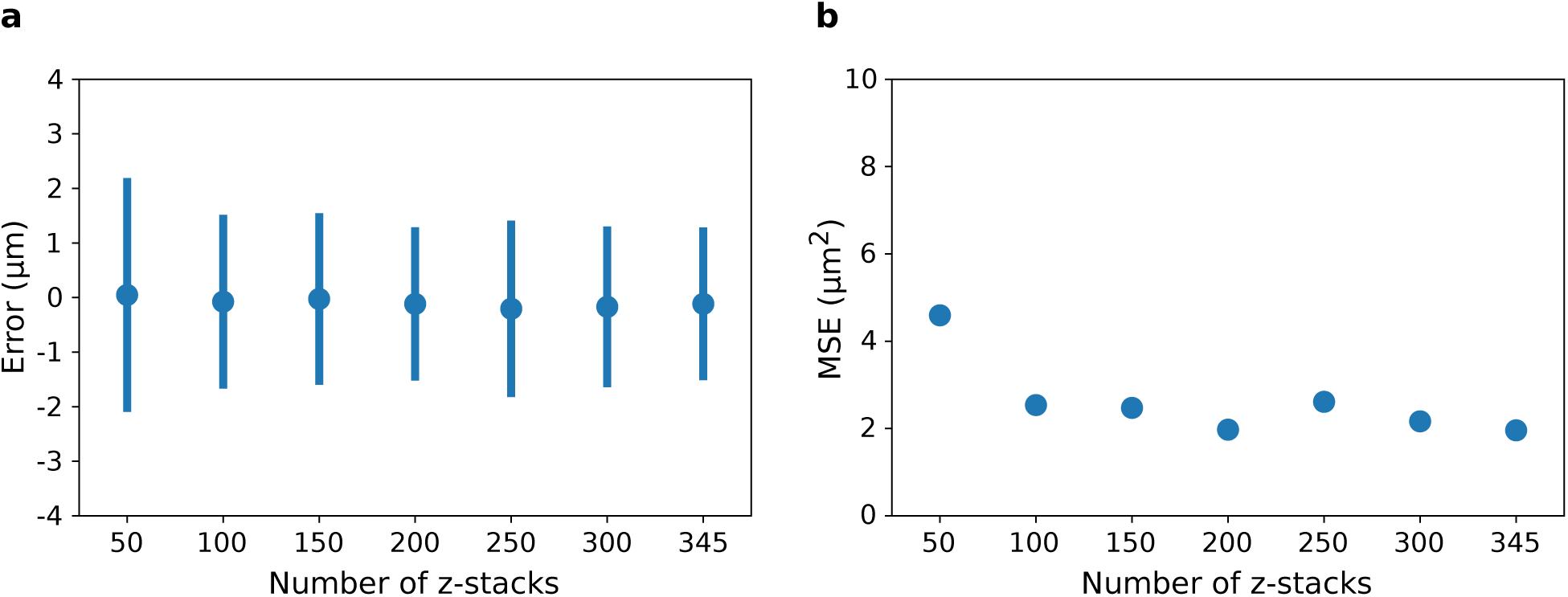
The relationship between accuracy and data set size. (a) Error as a function of the number of z-stacks in the data set. Symbols give the mean error and bars are ±1 standard deviation. (b) The mean squared error (MSE) vs. data set size.

### 2.5 Comparison of neural network and human focusing performance

To independently evaluate the performance of CNNs, we compared the accuracy of our autofocus CNN with four human annotators (H1–H4). We used for the computer (C) the best CNN model from the previous sections (4X downsampling, batch size 50, bin size 1, trained with 345 z-stacks). The evaluation was conducted using two separate tests: (1) identify the best in-focus image given a full z-stack, and (2) determine the numerical z-position of an individual image without using the z-stack for context. We provided both computer and human annotators with 100 z-stacks for part 1 and 100 individual images for part 2 that were randomly drawn from the testing data set.

As shown in Figure 5a, the CNN performed well in determining the correct in-focus image given an entire z-stack. Even though this was the easier of the two tests for the human annotators, since they could scan up and down in the z-stack to compare images, the CNN performance equaled the best human annotator and exceeded the other three. Moreover, in some cases, the human annotators showed a bias in determining the focal plane, while the computer showed minimum bias in the test.

**Figure 5:**
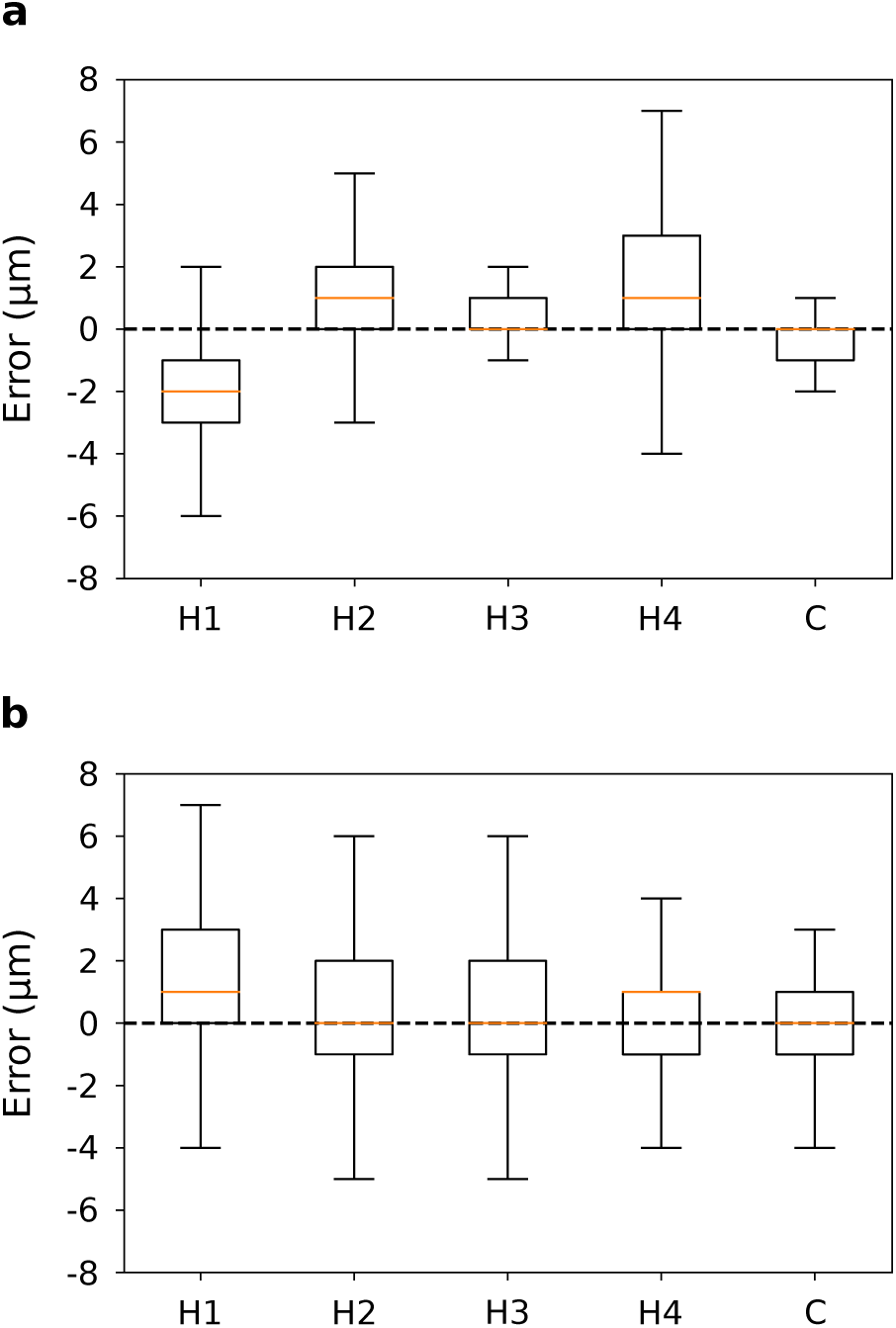
Comparison of the accuracy of the CNN and human annotators. (a) Error of four human annotators (H1-H4) compared with the CNN (C) in determining the in-focus image given a full z-stack to analyze. (b) Error in determining the numerical z-position of an image without using any additional context.

Figure 5b shows the results of determining the z-position of an individual image without access to nearby context. This was a very difficult test for the human annotators and they generally had a large degree of error. Here, the CNN performance exceeded that of all of the human annotators, being both more accurate and more precise. Interestingly, the CNN was more precise than the original human annotator who was one of the test subjects. If we assume that the annotator had a similar level of imprecision when annotating the training images, it appears that these errors average out during the training phase. Therefore, our trained CNN model was able to achieve accurate and consistent position inference that outperformed manual focusing.

### 2.6 Using the neural network to automatically maintain focus during live-cell imaging

Finally, we connected our best CNN model to the focal control of our microscope through a software interface. The CNN processes the live microscopy images at a rate of 20 Hz to estimate the current focal position of the sample. The controller then automatically adjusts the z-position of the objective in real-time to maintain the focal prediction within ±1 *μ*m of the focal plane. Figure 6 shows the result of a 24 hour growth experiment. Here, we purposely allowed the temperature of the environmental chamber to vary by 3°C to induce expansion effects on the stage (see Figure S2). The stage position varies by 40 *μ*m during the 24 hour period, but the sample is maintained in precisely the desired focal range.

**Figure 6:**
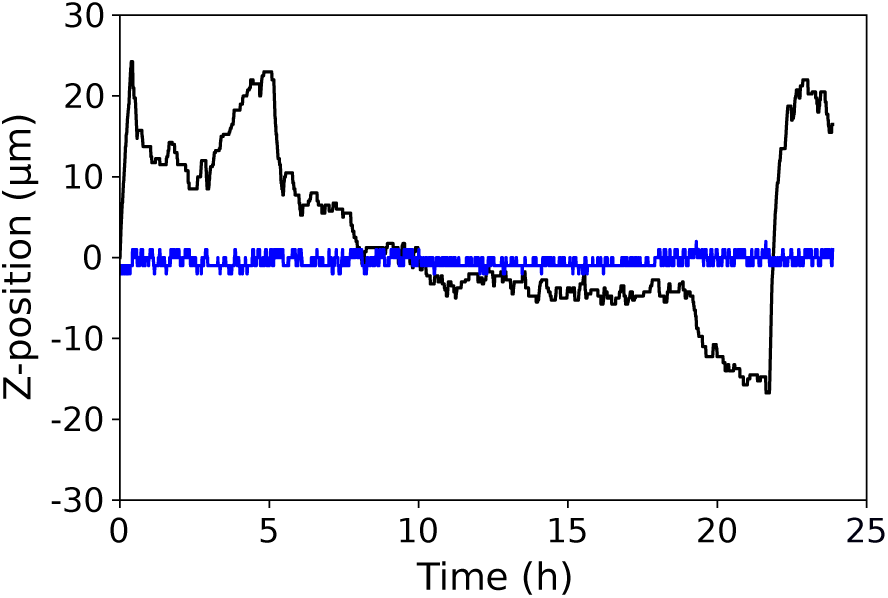
CNN controlled autofocusing during live-cell imaging of yeast. The (black) stage position and (blue) sample focal position over the course of a 24 hour growth experiment.

Moreover, focus is well-controlled regardless of large changes that occur in the field of view. Over the course of the 24 hour growth experiment the number of cells in the frame increases from fifteen to many hundreds (see Video S3). Additionally, because we did not affix the cells to the coverslip in this experiment, cells and cell clusters move around and occasionally drift into and out of frame. Focus is maintained within the set range regardless of these variations. The CNN is robust to these changes because it does not predict focus using any specific features but globally over the entire image.

## 3 Conclusion

Autofocus is of great importance in high-throughput microscopy of biological samples. It allows unattended imaging over large length and time scales. In this work, we presented a convolutional neural network (CNN) approach for inferring the focal position of cells under bright-field microscopy and for using the CNN output to reliably and accurately control a microscope in realtime to maintain focus. We have developed this method to assist in collecting high-throughput, live-cell imaging of yeast growing in a microfluidic device. Our method enables us to maintain focus on the cell monolayer during long-duration imaging and while rapidly scanning the stage to image the entire device, which may have height imperfections along its length.

In comparison with other autofocus approaches, our method does not require physical calibration nor the acquisition of z-stacks during imaging, and is robust to noise, optical artifacts, and features other than cells (e.g. structures in the microfluidic device). Even though the presented method provides some advantages over other forms of autofocus control, it is system specific in terms of microscope objective, imaging mode, and cell type. It is unlikely that a trained CNN model would work well if any of these variables were changed. Nevertheless, given that only a relatively small data set of 100 z-stacks is needed and an adequate CNN can be trained in a few days on a GPU, construction of system specific autofocus models is quite practical.

Additionally, it would be possible, given a large and varied data set and a deeper neural network, to train a single model to recognize focal position in many different systems. Our method therefore provides a novel conceptual and practical framework for automated focus control in high-throughput imaging of biological samples, and we anticipate that this framework is generally applicable in the field of light microscopy where unattended and real-time control of focal position is desired.

Finally, while our CNN model only infers the best single focal position for an entire image, by using deconvolution techniques [36, 37] the focal position of all individual cells within the image could also be obtained. Then, all cells within the field of view could be focused on and imaged individually or complex spatially varying features could be tracked and followed in the z dimension. Such a method would assist in automatic high-throughput imaging of multiple cell layers within complex biofilms or tissues.

## 4 Materials and Methods

### 4.1 Cell growth and microscopy

The Yeast GFP Clone Collection STE20/YHL007C fusion strain (courtesy of Dr. John Kim) was used in this study. Yeast cells were initially cultured overnight in low-fluorescence synthetic defined media without histidine (SD-H) at 30°C. Cells were then loaded into a custom soft-lithography microfluidic device with a 6 *μ*m tall growth chamber containing 10 *μ*m diameter pillars on a 50 *μ*m grid to prevent chamber collapse. The device was fabricated according to standard protocols. Yeast cells were continuously supplied with fresh SD-H media through the entire imaging process.

Images were acquired using an Axiovert 35 inverted microscope (Zeiss) equipped with a custom automated stage and environmental chamber. A Fluar 40X oil-immersion objective with 1.30 NA (Zeiss) was used during all microscopy. Images were captured using a Flea3 FL3-U3-32S2M-CS camera (FLIR Systems) operating at 20 fps, 10 dB gain, and 50 ms exposure time. The image size was 2080×1552 pixels with a resolution of ~60 nm per pixel. The microscope, stage, camera, and light source were controlled by custom software. Z-stacks were acquired at room temperature and live-cell imaging was conducted at 30°C.

### 4.2 Neural network implementation

The neural network was implemented using the TensorFlow framework [38]. In TensorFlow, computation, shared state, and operations that mutate that state are represented by dataflow graphs. Specifically, functional operators, such as matrix multiplication and convolution, as well as mutable state and operations that update the state, are represented as nodes, while multi-dimensional arrays (tensors) that are input to or output from nodes are represented as edges. All training runs were executed on 4GB Tesla K20m GPUs (Nvidia). The code for training the neural network and inferring the focal position of images is given in Supplementary File S4. All training and testing data, along with the final neural network model, is available for download from our website (http://www.robertslabjhu.info/home/data/).

## Acknowledgments

The authors would like to thank the members of Roberts lab for discussions and for participating in the annotation tests. This work was supported by the National Science Foundation under grant number PHY-1707961.

## Author Contributions

E.R. designed the study, developed the software tool, and guided the project. L.W. and E.R. performed the experiments. E.R. and L.W. analyzed the data. E.R. and L.W. wrote the manuscript.

